# Long-read sequence capture of the hemoglobin gene clusters across species

**DOI:** 10.1101/297796

**Authors:** Siv Nam Khang Hoff, Helle T. Baalsrud, Ave Tooming-Klunderud, Morten Skage, Todd Richmond, Gregor Obernosterer, Reza Shirzadi, Ole Kristian Tørresen, Kjetill S. Jakobsen, Sissel Jentoft

## Abstract

Combining high-throughput sequencing with targeted sequence capture has become an attractive tool to study specific genomic regions of interest. Most studies have so far focused on the exome using short-read technology. These approaches are not designed to capture intergenic regions needed to reconstruct genomic organization, including regulatory regions and gene synteny. Here, we demonstrate the power of combining targeted sequence capture with long-read sequencing technology for comparative genomic analyses of the hemoglobin (Hb) gene clusters across eight species separated by up to 70 million years. Guided by the reference genome assembly of the Atlantic cod (*Gadus morhua*) together with genome information from draft assemblies of selected codfishes, we designed probes covering the two Hb gene clusters. Use of custom-made barcodes combined with PacBio RSII sequencing led to highly continuous assemblies of the LA (~100kb) and MN (~200kb) clusters, which include syntenic regions of coding and intergenic sequences. Our results revealed an overall conserved genetic organization and synteny of the *Hb* genes within this lineage, yet with several, lineage-specific gene duplications. Moreover, for some of the species examined, we identified amino acid substitutions at two sites in the *Hbb1* gene as well as length polymorphisms in its regulatory region, which has previously been linked to temperature adaptation in Atlantic cod populations. This study highlights the use of targeted long-read capture as a versatile approach for comparative genomic studies by generation of a cross-species genomic resource elucidating the evolutionary history of the *Hb* gene family across the highly divergent group of codfishes.

## Introduction

The rapid advancement of high-throughput sequencing has over the last decade revolutionized genomic research with the increasing numbers of whole genome resources available for multiple vertebrate species, including the diverse group of teleost fishes (Volff 2004; Ellegren 2014; Goodwin *et al*. 2016; Malmstrøm *et al*. 2016; 2017). However, whole genome sequencing (WGS) and generation of high quality genome assemblies are still considered costly and time-consuming (Jones & Good 2015). For investigations concerning specific genomic regions there is no need for complete genome information, which has spurred the development of reduction complexity approaches such as targeted sequence capture (Teer *et al*. 2010; Grover *et al*. 2012; Samorodnitsky *et al*. 2015). The basic idea of targeted sequence capture involves design of specific probes covering the particular genomic area of interest generating an enriched coverage of the targeted sequences (Turner *et al*. 2009; Grover *et al*. 2012). Most studies using targeted sequence capture have to a large extent been directed towards the exome, often supported by the existence of a reference genome (Broeckx *et al*. 2014; Yoshihara *et al*. 2016), or transcriptome assemblies (Syring *et al*. 2016). Recent reports have, however, been focusing on off-target sequences in noncoding regions (Guo *et al*. 2012; Syring *et al*. 2016; Yoshihara *et al*. 2016; Morin *et al*. 2016), as they may contain crucial regulatory elements varying in sequence and length between populations or species and could be of functional and evolutionary importance (Woolfe *et al*. 2004).

To date, comparative studies using sequence capture have been mainly based on short-read sequencing technology and probe design targeting genic regions (George *et al*. 2011; Samorodnitsky *et al*. 2015; Bragg *et al*. 2015; Li *et al*. 2015). Consequently, construction of continuous sequences enabling resolution of gene organization across species has not yet been looked into. Comparative genetic studies of gene organization or synteny requires longer, more continuous stretches of DNA containing more than one gene (Huddleston *et al*. 2014). By its ability to span long stretches of repeats, long-read sequencing technology has been successfully applied to improve genome assembly statistics and generation of highly continuous genome assemblies for a growing number of species (English *et al*. 2012; Kim *et al*. 2014; Tørresen *et al*. 2017; Kolmogorov *et al*. 2018; Tørresen *et al*. 2018). Incorporation of long PacBio reads have for instance resulted in a significantly improved version of the Atlantic cod (*Gadus morhua*) genome assembly, i.e. a 50-fold increase in sequence continuity and a 15-fold reduction in the proportion of gap (Tørresen *et al*. 2017). Furthermore, recent studies report the combination of capture and long read sequencing as highly efficient in enriching and assembly of full length complex genes as well as detailed characterization of chromosomal structural variations (Wang *et al*. 2015; Witek *et al*. 2016; Giolai *et al*. 2017). Correspondingly, utilizing long-read sequencing technology in combination with targeted capture could yield longer continuous assemblies of specific genomic regions of interest, allowing in-depth comparative genetic studies including synteny analyses, in species where reference genomes are not available.

In fishes, the hemoglobin (*Hb*) gene family, encoding the protein subunits Hba and Hbb, has shown to be of importance for ecological adaptation, as environmental factors such as temperature directly influences the ability of Hb to bind O2 at respiratory surfaces and its subsequent release to tissues (Wells 2005). In a recent report, a characterization of the *Hb* gene repertoire by comparative draft genome analysis uncovered a remarkably high *Hb* gene copy variation within the codfishes (Baalsrud *et al*. 2017). Based on the gene copy number a negative correlation between the number of *Hb* genes and depth of which the species occur was observed, as well as signs of diversifying selection on the gene paralogues suggesting that the variable environment in epipelagic waters have facilitated a larger more diverse *Hb* gene repertoire (Baalsrud *et al*. 2017). However, the rather fragmented draft genomes did not allow for reconstruction of the gene synteny and a deeper understanding of the evolution of the Hb-clusters.

Moreover, two tightly linked polymorphisms at amino acid positions 55 and 62 of the Hbb1-globin suggested associated with thermal adaptation are demonstrated in Atlantic cod populations. These polymorphisms exhibit a latitudinal cline in allele frequency in populations inhabiting varying temperature and oxygen regimes for Atlantic cod in the North Atlantic and Baltic Sea (Andersen *et al*. 2009). Populations found in the southern regions display the Hbb1-1 variant (Met55Lys62), whereas more northern populations largely display the Hbb1-2 variant (Val55Ala62) (Andersen *et al*. 2009). The Hbb1-1 variant has been shown to be insensitive to temperature whereas Hbb1-2 is temperature dependent with a higher O2 affinity than Hbb1-1 at colder temperatures (Andersen *et al*. 2009), however, this has been questioned by Barlow *et al*. 2017. Additionally, an indel polymorphism within the promoter of the *Hbb1* gene has been reported to be in linkage disequilibrium with the above-mentioned polymorphisms (Star *et al*. 2011). Examination of multiple Atlantic cod populations uncovered that a longer promoter variant is associated with *Hbb1-2* and found to up-regulate its gene expression at higher temperatures, i.e. aiding in the maintenance of total oxygen-carrying capacity (Star *et al*. 2011).

In teleosts, the *Hb* genes are found to reside at two distinct genomic regions, the MN and LA cluster. Earlier reports have shown that there is a high evolutionary turnover of *Hb* genes across teleosts, with lineage-specific duplications and losses, which is in stark contrast to genes flanking the *Hb* genes, where the synteny is highly conserved (Quinn *et al*. 2010; Opazo *et al*. 2012; Feng *et al*. 2014). In this study, the overall goal was to elucidate the evolutionary past of the *Hb* clusters – including *Hb* genes, flanking genes and intergenic sequences – within the phylogenetically diverse group of codfishes (Gadiformes) by taking advantage of long read sequencing technology combined with targeted sequence capture. Eight codfish species were carefully selected on the basis of both phylogenetic and habitat divergence, implying that they are exposed to a variety of environmental factors as well as displaying distinct life-history traits. A highly continuous genome assembly of Atlantic cod (Tørresen *et al*. 2017) as well as low coverage draft genome assemblies of all eight species (Malmstrøm *et al*. 2017) were used in the design of the probes covering the genomic regions of interest. To enable targeted sequence capture for PacBio RSII sequencing, we modified the standard protocol for sequence capture offered by NimbleGen, i.e. the SeqCap EZ (Roche NimbleGen), as well as generating custom-made barcodes. This combined approach resulted in successful capturing and assembling of the two *Hb* gene clusters across the codfishes examined. The generation of highly continuous assemblies – for most of the species – enabled reconstruction of micro-synteny revealing lineage-specific gene duplications and identification of a relatively large and inter-species variable indel located in the promoter region between the *Hbb1* and *Hba1* genes.

Our study demonstrates that combining sequence capture technology with long-read sequencing is a highly efficient and versatile method to investigate specific genomic regions of interest – with respect to micro-synteny, regulatory regions and genetic organization – across distantly related species where genome sequences are lacking.

## Results

### Capture and *de novo* assembly of the target regions

The probe design (workflow schematically shown in Figure 1) resulted in a total of 7057 capture probes based on the target region in Atlantic cod, covering 337 kbp of sequence. 26774 probes were designed for the additional codfishes, covering a total of 1.82 Mbp of target sequence. The target region and the *Hb* gene clusters were successfully captured and enriched for eight codfishes; Atlantic cod (*Gadus morhua*), haddock (*Melanogrammus aeglefinus*), silvery pout (*Gadiculus argenteus*), cusk (*Brosme brosme*), burbot (*Lota lota*), European hake (*Merluccius merluccius*), marbled moray cod (*Muraenolepus marmoratus*), and roughhead grenadier (*Macrourus berglax*), with number of reads spanning from 35573 to 73005 (Table 1). The average read length was 3032 bp, varying from 2836 bp in European hake to 3265 bp in burbot, resulting in the capture of an average of 16.71 Mbp per species (Table 1). By mapping reads back to the capture target region we found that the average mapping depth was variable across the target region for all species (Figure 2 and 3). Because of the skewed distribution of mapping depth, we also calculated median depth, which was, as expected, the highest for Atlantic cod at 242x (Table S1). The median mapping depth was consistently high for most of the other species as well, with the lowest for roughhead grenadier (12x). Both median and average depths for the MN region were persistently higher than for the LA region for all species, with the exception of silvery pout (Table S1). Furthermore, positions with high degree of mapping corresponded to the location of the genes used in the design of the capture probes across all species (Figure 2 and Figure 3). The percentage of reads mapping to the target region ranged from 25-43%, however, the percentage of the target region covered by reads ranged from 53-100% with five species having more than 90% of the target region covered by reads (Figure 4c and Table S1).

**Table 1.** Number of reads and bases captured and sequenced for each species, and number of utgs, largest utg and N50 in the assemblies.

| Species | Latin name | Number of reads | Number of bases | Number of utgs | Largest utg (bp) | N50 (bp) |
| --- | --- | --- | --- | --- | --- | --- |
| **Atlantic cod** | *Gadus morhua* | 73005 | 217252583 | 278 | 79 020 | 7 728 |
| **Haddock** | *Melanogrammus aeglefinus* | 35573 | 107839552 | 227 | 52 433 | 7 227 |
| **Silvery pout** | *Gadiculus argenteus* | 69775 | 212519845 | 410 | 35 801 | 7 098 |
| **Cusk** | *Brosme brosme* | 55348 | 175883008 | 394 | 64 145 | 7 322 |
| **Burbot** | *Lota lota* | 56155 | 165360828 | 205 | 70 602 | 8 055 |
| **European hake** | *Merluccius merluccius* | 65661 | 180558336 | 311 | 31 558 | 6 523 |
| **Marbled moray cod** | *Muraenolepus marmoratus* | 52076 | 148100933 | 455 | 30 019 | 6 632 |
| **Roughhead grenadier** | *Macrourus berglax* | 46195 | 129085001 | 325 | 35 216 | 7122 |

**Figure 1:**
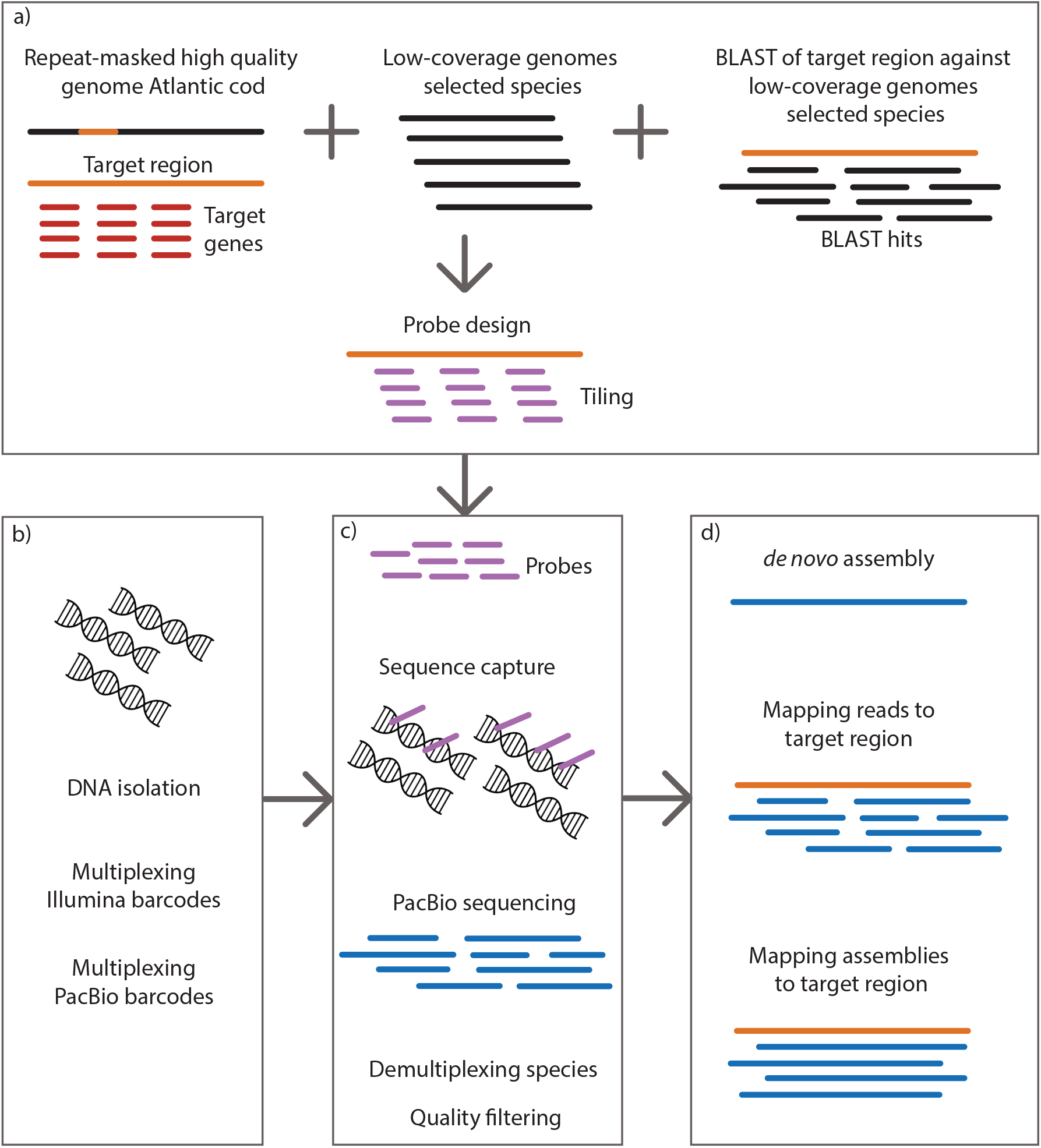
Flowchart of sequence capture approach. a) Sequence data from the Atlantic cod genome (Tørresen *et al*. 2017; gadMor2) combined with gene sequences of target genes and sequences from low coverage genomes of the additional codfishes are combined to generate probes. b) Isolated DNA is multiplexed with Illumina and PacBio barcodes. c) Raw reads for each species are used to score all probes, ensuring that no repeated sequences are present. DNA Probes are used in solution on isolated DNA for all of the included species, hybridizing to the target sequences. Target sequences are then captures and sequences on the PacBio RSII sequencing platform. d) Downstream bioinformatics includes de-multiplexing of reads and trimming, making the reads ready for downstream analysis such as mapping and *de novo* assembly.

**Figure 2:**
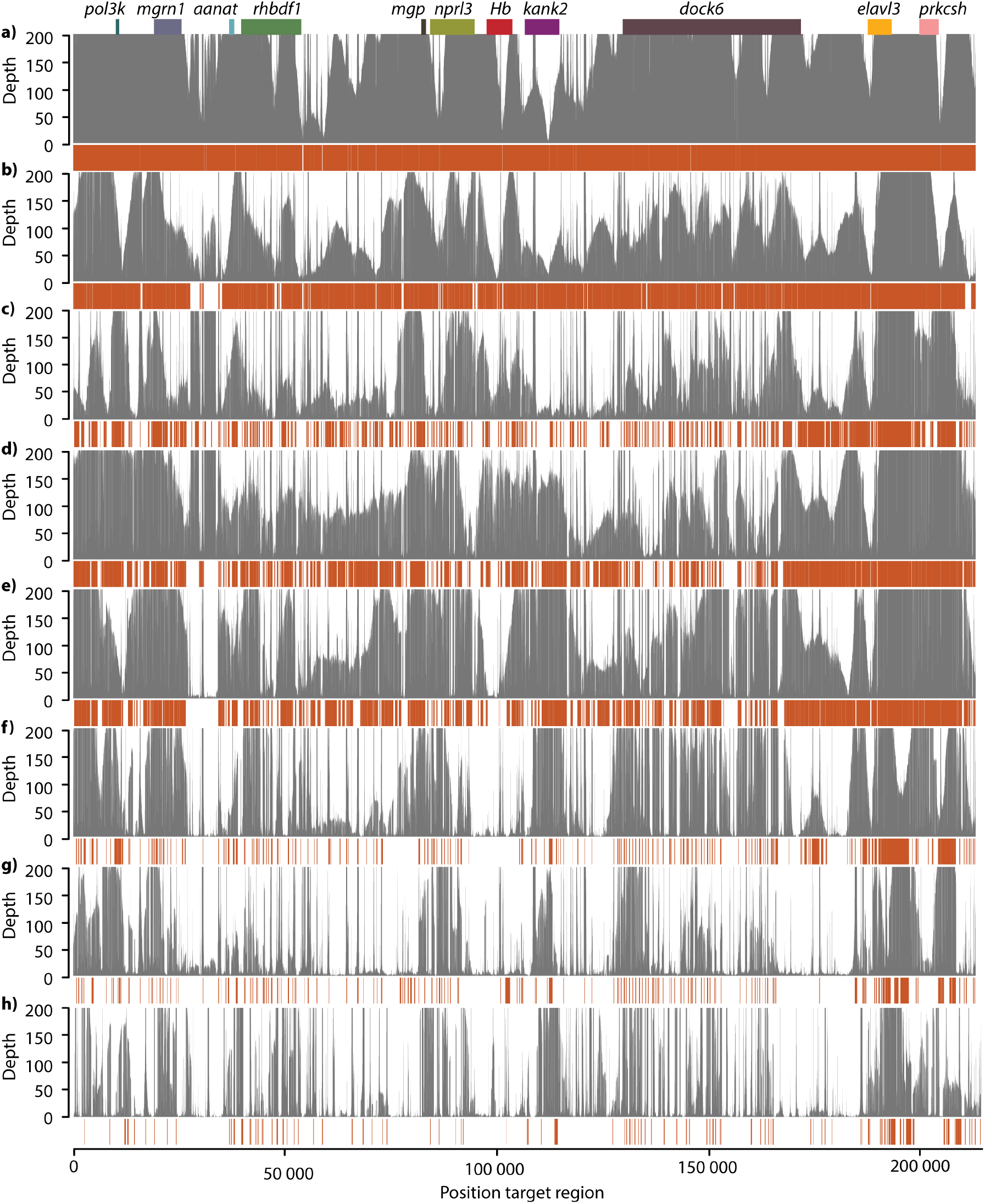
Mapping of reads and assemblies against the MN target region. Each panel shows the reads and *de novo* assembly mapped against the MN target region in grey and orange, respectively, for species a.) Atlantic cod, b) haddock, c) silvery pout, d) cusk, e) burbot, f) European hake, g) marbled moray cod and h) roughhead grenadier. The positions of genes in the target region are indicated at the top.

**Figure 3:**
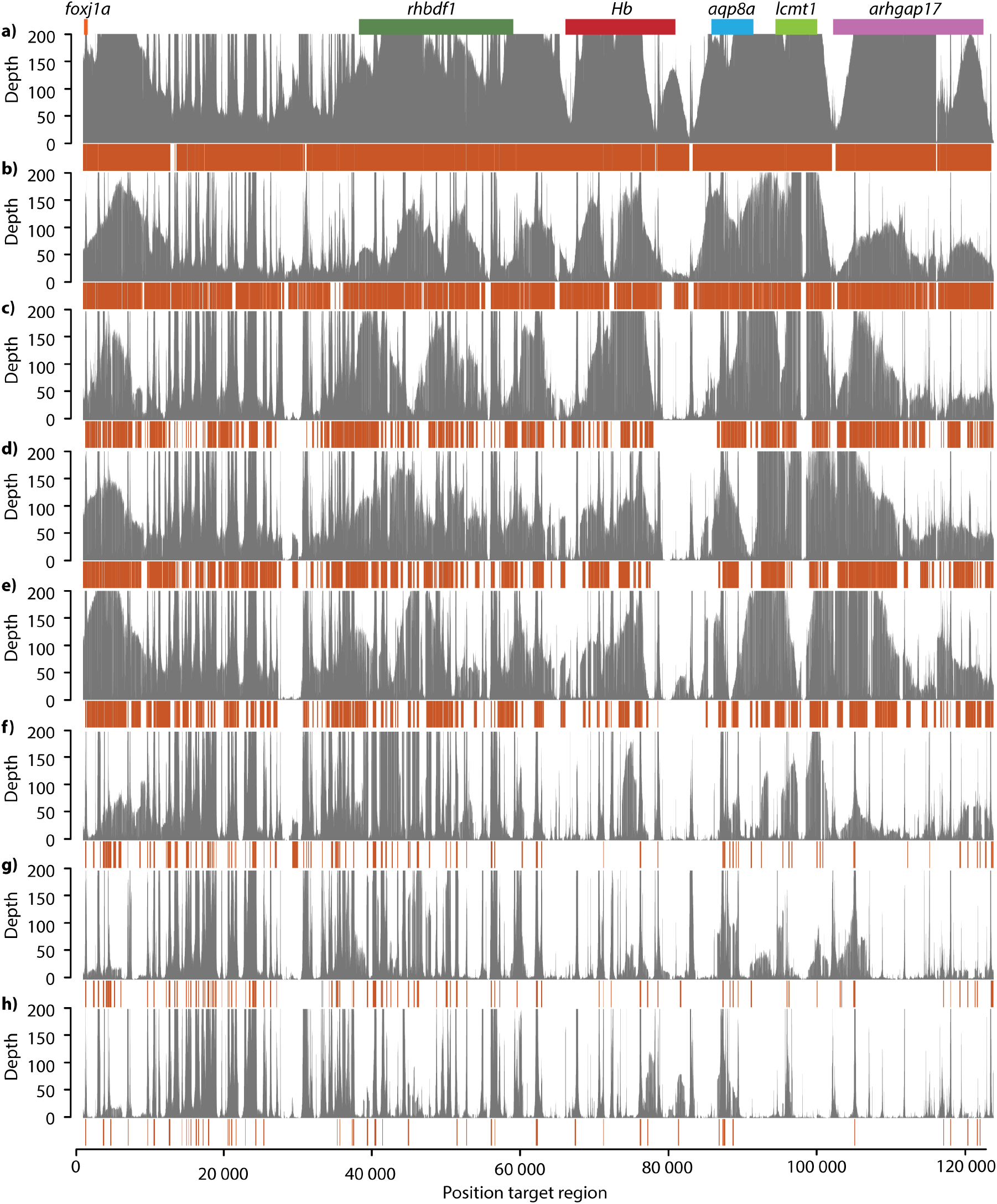
Mapping of reads and assemblies against the LA target region. Each panel shows the reads and *de novo* assembly mapped against the LA target region in grey and orange, respectively, for species a) Atlantic cod, b) haddock, c) silvery pout, d) cusk, e) burbot, f) European hake, g) marbled moray cod and h) roughhead grenadier. The positions of genes in the target region are indicated at the top.

**Figure 4:**
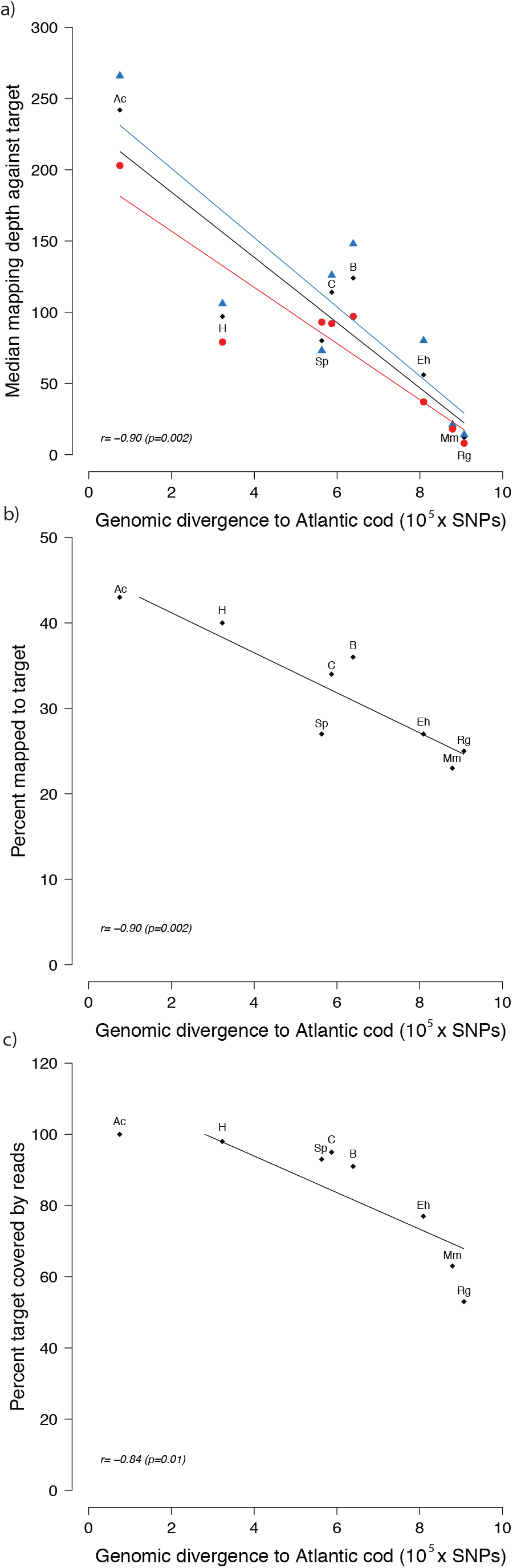
The relationship between capture success and genomic divergence to Atlantic cod. Linear regression of the relationship between the genomic divergence to Atlantic cod (SNPs × 10^5^) and a) median mapping depth for the MN region (blue), LA region (red) and the combined target region (black); b) the percentage of reads mapping to the target region; c) the percentage of the target region covered by reads to a minimum depth of 10x. For each regression the correlation coefficient, r, is shown along with a p-value. Each data point is labeled by species according to this code: Ac=Atlantic cod, H=haddock, Sp=silvery pout, C=cusk, B=burbot, Eh=European hake, Mm=marbled moray cod and Rg=roughhead grenadier.

To address factors influencing capture success we compared various capture statistics to overall genomic divergence between the Atlantic cod genome and independent WGS data for each species from (Malmstrøm *et al*. 2017) (Table S1). We found a strong negative correlation between genomic divergence to Atlantic cod and median mapping depth against the target region (r=-0.90, Figure 4a), percent of reads mapped to the target region (r=-0.90, Figure 4b), and percentage of reads mapped to the target region (r=-0.84, Figure 4c).

We constructed *de novo* assemblies with quite consistent assembly statistics across species. Contig N50 ranged from 8055 bp in burbot to 6523 bp in European hake and the total number of contigs varied from 205 in burbot to 455 in marbled moray cod. However, there was some variation in the size of the largest contig, which ranged from 79 kbp in Atlantic cod to 30 kbp in marbled moray cod (Table 1). To evaluate whether the assemblies represent the actual target regions we mapped the *de novo* assemblies for each species to the target region in Atlantic cod, for which the capture design is largely based upon (Figure 2 and 3). As expected, the assemblies corresponded to the regions with high coverage of reads, i.e. the areas of the target region containing genes included in the probe design.

### Synteny of the *Hb* gene regions

Our capture design combined with long-read PacBio sequencing allowed us to reconstruct micro-synteny of the MN and LA regions for Atlantic cod, haddock, silvery pout, cusk, burbot, European hake, marbled moray cod and roughhead grenadier (Figure 5). From the *de novo* assemblies, we were able to identify the majority of the *Hb* genes and all of the flanking genes, which show that our capture design was successful. However, the degree of continuity varied in the different assemblies. In Atlantic cod, haddock, silvery pout, cusk, burbot and European hake we could infer micro-synteny revealing that *Hb* and their flanking genes organization largely followed what has previously been reported for Atlantic cod (Figure 5) (Star *et al*. 2011). We found *Hbb4* only to be present in Atlantic cod (Figure 5b), which is in line with Baalsrud *et al*. 2017. Furthermore, the *de novo* assemblies confirmed a linage-specific duplication of *Hbb2* in the roughhead grenadier (Baalsrud *et al*. 2017). Additionally, we identified a complete Hba4-like gene in the assembly of the marbled moray cod, not earlier identified in this species. However, the *Hba4-like* gene in marbled moray cod is likely a pseudogene due to a frameshift mutation causing multiple stop codons. Furthermore, we were able to identify most of the *Hb* genes reported in the recent study by Baalsrud *et al*. 2017, however, a few are missing from our dataset (Figure 5a and b). Pairwise sequence alignment of these paralogous *Hb* genes from Baalsrud *et al*. 2017 revealed sequence identities up to 98 % (Table S2).

**Figure 5:**
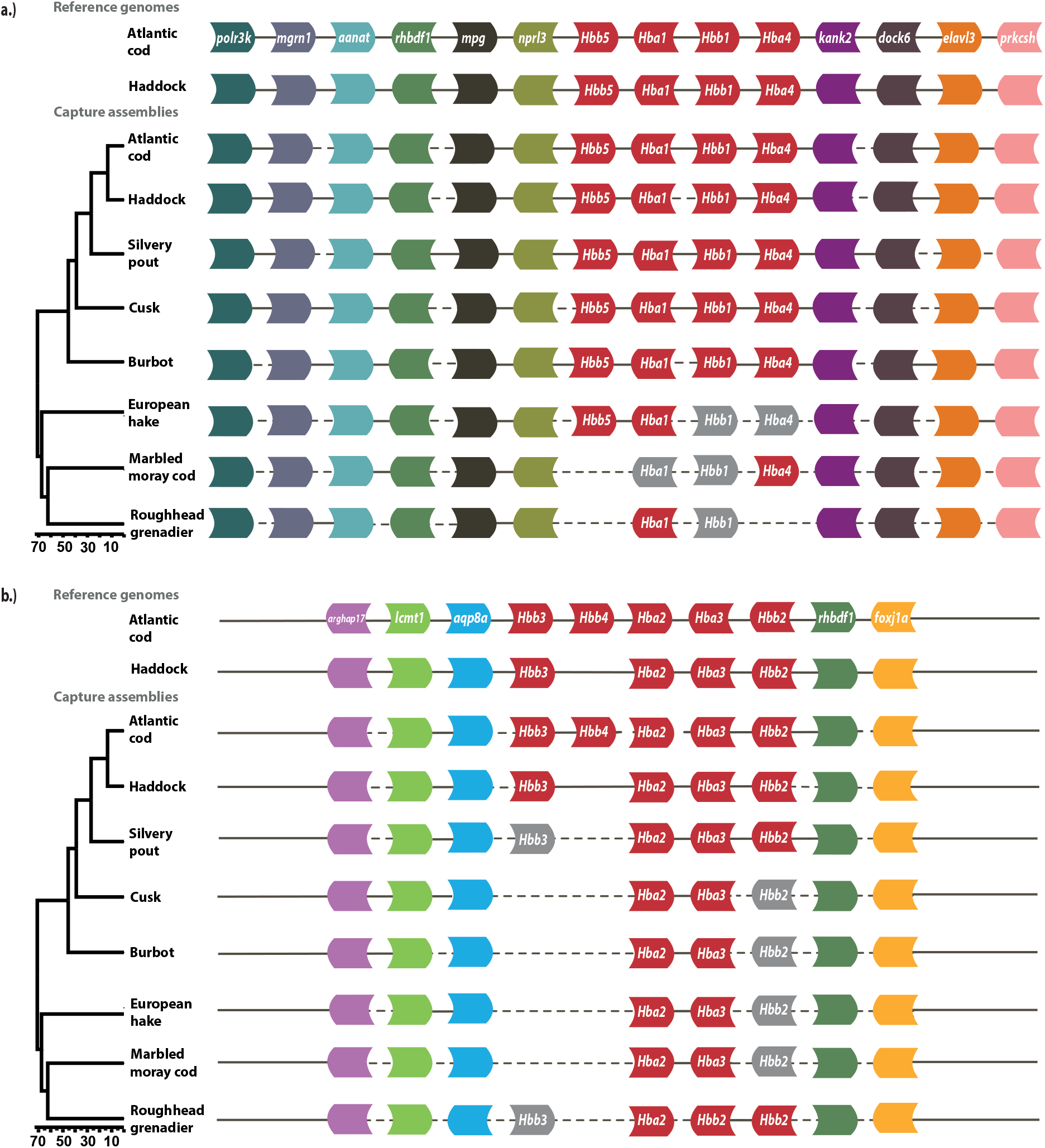
Synteny of the Hb gene clusters. Genomic synteny of the hemoglobin gene clusters shown at the top for the genomes of Atlantic cod (Tørresen *et al*. 2017; gadMor2) and haddock (Tørresen *et al*. 2018; melAeg). Below, the genomic synteny inferred from the *de novo* assemblies for all of the species included in the capture experiment. Stippled lines indicate assembly gaps – here we assume that the orientation of genes corresponds to the genomes of Atlantic cod and haddock. Gray boxes indicate genes that have been identified in Baalsrud *et al*. (2017) but are absent in the *de novo* assemblies. a) Synteny across the MN region b) Synteny across the LA region.

### Target region in the haddock and Atlantic cod genome assemblies

As a proof of concept, we reconstructed synteny of the target region in the most recent genome assemblies of Atlantic cod (Tørresen *et al*. 2017; gadMor2) and haddock (Tørresen *et al*. 2018; melAeg). In Atlantic cod, the MN region is located on linkage group 2 (Figure 5a) and LA on linkage group 18 (Figure 5b), in haddock MN is located on scaffold MeA_20160214_scaffold_771 (Figure 5a) and LA on scaffold MeA_20160214_scaffold_1676 (Figure 5b). The overall synteny in Atlantic cod was congruent with Wetten *et al*. 2010 except for the relative direction of the genes *foxj1a* and *rhbdf1*. Furthermore, the organization of *Hbs* and their flanking genes in the genome assembly of haddock is conserved compared to Atlantic cod with the exception of *Hbb4* in the MN region, which is absent in haddock (Figure 5).

### Repetitive sequences in the in the *Hb* gene regions

Quantifying the amount of repetitive sequences in the target region(s) was only possible for Atlantic cod (gadMor2) and haddock (melAeg), for which high-quality genome assemblies exist. The amount of repetitive sequences in the target region differed between the MN cluster and the LA cluster in Atlantic cod. The MN region (214 kb) contained a total of 10.7% repeated sequences, including 1.0% retro-elements, 1.3% transposons, 5.8% simple repeats, and 2.6% of various low complexity and unclassified repeated sequences (Table S3). In comparison, in the smaller LA region (123 kb) the proportion of repeated sequences was twice as high (20.3%), which comprised of 2.8% retro-elements, 2.4% transposons, 13.8% simple repeats, and 1.3% of various low complexity and unclassified repeated sequences. Furthermore, the orthologous target regions in haddock followed the same pattern. The MN region contained 16.3 % repeated sequences, in contrast to 19.8 % found in the LA region (Table S3).

### Insertions and deletions in the promoter region of *Hbal* – *Hbb1*

The previously shown 73 bp indel in the bi-directional promoter region of *Hba1* and *Hbb1* – discerning the cold-adapted migratory Northeast Artic (NEA) cod from the more temperate-adapted southern Norwegian coastal (NC) cod (Star *et al*. 2011) – was confirmed by the improved version of the NEA cod assembly (gadMor2). The continuity of our capture assemblies (Figure 5) enabled location of the orthologous captured regions in haddock, silvery pout and cusk. In each of the species an indel of variable length were identified (Figure 6). Compared to the long promoter variant – found to be linked with the *Hbb1-2* in Atlantic cod – the indel is shorter in the other species by 11 bp in haddock, 22 bp in silvery pout and 56 bp in cusk (Figure 6). Although the indels are varying in length, the conserved flanking sequences in the alignment clearly show that they represent orthologous regions. Moreover, we found the amino acid positions at 55 and 62 of the *Hbb1* gene to vary between species; Haddock has Val55-Lys62, silvery pout has Met55-Gln62, while cusk has Met55-Lys62 similarly to NEA cod (Figure 6). Additionally, we investigated amino acid positions 55 and 62 in the *Hbb1* gene across a number additional codfish species for which we have available gene sequences from Baalsrud *et al*. 2017, revealing these sites to be variable across this lineage (Table S4). Ancestral reconstruction of *Hbb1* demonstrated that the ancestral state in position 55 was Met in codfishes, and in position 62 was Lys in all codfishes except *Bregmaceros cantori* (Supplementary Figures S1 and S2).

**Figure 6:**
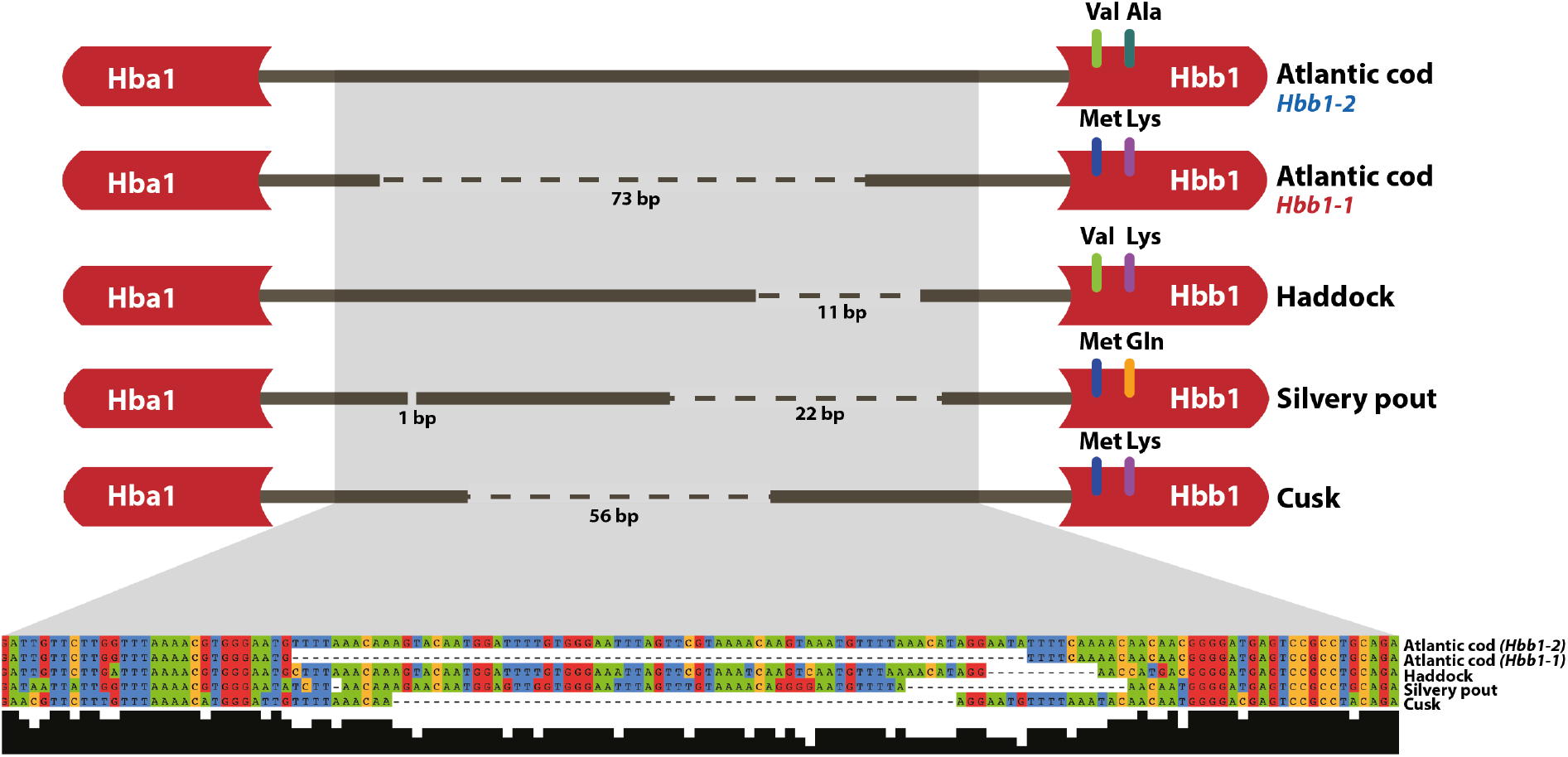
Polymorphisms in the bi-directional promoter between *Hba1* and *Hbb1* for five species in the Gadidae family. A schematic representation of *Hba1* and *Hbb1* with the promoter region between them. The region contains an indel polymorphism of variable length across the five species, as indicated by gaps. For each species/variant the alignment is shown along with amino acid substitutions at positions 55 and 62 in the translated part of the *Hbb1* gene.

## Discussion

### Capture of *Hb* gene clusters with 70 million years divergence time reveal conserved synteny and lineage-specific *Hb* duplications

We here demonstrate a successful in-solution targeted sequence capture and assembling of coding and noncoding sequences of the *Hb* clusters from codfish species separated by up to 70 million years (My) of evolution. Two features make our approach unique from earlier studies. First, the target regions consisted of both coding and noncoding genomic sequences. Second, we designed custom-made probes in order to utilize the long-read PacBio sequencing platform. In contrast to previous targeted capture sequencing studies based on short-read sequencing technologies (George *et al*. 2011; Mascher *et al*. 2013) our approach enabled the generation of highly continuous assemblies of the *Hb* clusters across distantly related codfishes.

The organization and orientation of the *Hb* flanking genes that we identified were conserved across all species (Figure 5a and b). However, in concordance with earlier studies of the *Hb* region, we found significant variation in copy numbers of the *Hb* genes, with linage specific duplications and losses (Star *et al*. 2011; Opazo *et al*. 2012; Feng *et al*. 2014; Baalsrud *et al*. 2017). We only found *Hbb4* in Atlantic cod, supporting earlier studies showing that *Hbb4* is the result of a recent duplication in this species (Borza *et al*. 2009; Baalsrud *et al*. 2017). Interestingly, the presence of two copies of *Hbb2* on the same contig in the roughhead grenadier *de novo* assembly confirmed a lineage-specific gene duplication of *Hbb2* in this species. This duplication was postulated in recent study of *Hbs* in codfishes (Baalsrud *et al*. 2017), but due to the lack of synteny in the draft genome assemblies, it was not possible to determine with certainty. Additionally, a copy of the *Hba4* was found in the *de novo* assembly of the marbled moray cod not found in the previous study by Baalsrud *et al*. (2017). The presence of a frame-shifting mutation that is causing multiple stop codons indicated that this *Hba4* gene is most likely a pseudogene. *Hba4* is also a pseudogene in the closely related species *Mora moro, Trachyrincus scabrus, T. murrayi* and *Melanonus zugmayeri* (Baalsrud *et al*. 2017). Although we identified most of the *Hb* genes from Baalsrud *et al*. (2017), a few were absent from this dataset (Figure 5a and b), which we suspect may be due to collapse of paralogous *Hb* genes, as they may have as high as 98% sequence identity (Table S2).

### Length variation in the bi-directional *Hbal-Hbbl* promoter within the codfishes

The discovery of a promoter of variable length between *Hba1* and *Hbb1* in different species (Figure 6) was concordant with earlier findings of length variation in the homologous region in different populations of Atlantic cod (Star *et al*. 2011). The migratory NEA cod population has been shown to harbor the 73 bp longer variant at a higher frequency compared to coastal cod populations (see Figure 6 and Star *et al*. 2011). Interestingly, we found relatively long promoters with high sequence similarity to the NEA cod indel in haddock and silvery pout. In contrast, cusk displayed a relatively short promoter, however, still 17 bp longer than in NC cod (Figure 6). Furthermore, we found the amino acid positions 55 and 62 in *Hbb1*, known to be polymorphic in Atlantic cod, to be variable across all codfishes included in this study (Figure 6). Investigations of the same positions in a number additional codfishes for which we have available gene sequences (Baalsrud *et al*. 2017), revealed that these positions are highly variable across this linage (Table S4). Notably, the most likely ancestral state of codfish *Hbb1* is Met55Lys62 (Supplementary Figures S1 and S2). Cusk and the coastal/southern Atlantic cod thus both display the ancestral state as well as a short promoter, although the cusk promoter was 17 bp longer (Figure 6). Collectively, these results suggest two different scenarios for promoter length evolution. Scenario 1: The short promoter represents the ancestral state of the Gadidae-family (including cusk and Atlantic cod; see Malmstrøm *et al*. 2016) and that silvery pout and some populations of Atlantic cod have evolved a longer promoter. Scenario 2: The long promoter is the ancestral state with independent deletions of variable lengths in cusk, silvery pout, haddock and coastal/southern Atlantic cod (*Hbb1-1*). To disentangle this, we would need to obtain promoter sequences from additional gadiform species. Regardless, the short-long promoter polymorphism has been maintained throughout speciation events based on the presence of both variants in Atlantic cod. Moreover, in both scenarios, cusk and coastal/southern Atlantic cod (*Hbb1-1*) have maintained the ancestral Met55Lys62, while silvery pout, haddock and NEA cod (*Hbb1-2*) have acquired substitutions at these positions due to similar selection pressures or genetic drift. In this regard, it could be mentioned that the NEA cod, haddock and silvery pout display migratory behavior (e.g. diurnally feeding movements as well as seasonal spawning migrations) compared to the more stationary cusk and coastal cod (Eschemeyer & Fricke 2017) which could mean that they have a higher O2 demand and are exposed to greater temperature variation, which in turn has selected for a temperature-dependent long promoter. Furthermore, given that promoter length and positions 55/62 at *Hbb1* are important genetic components of temperature adaptation in Atlantic cod populations (Star *et al*. 2011), they most likely play a role in temperature adaptation in the other codfishes.

### Assembly success affected by probe design and repeat content

In some species, nearly the complete target region is assembled in large contigs containing multiple genes including cusk, whereas in other species such as the more distantly related roughhead grenadier, the cluster is more fragmented (Figure 5). In all species, the areas of the target regions that harbor genes of which probes are designed for, as well as any areas containing repeated sequences, have very high depths in comparison to the areas of intergenic sequences (Figure 2 and 3). This poses a challenge for the assembly software, which is based in the assumption of uniform depth over the sequencing data (Miller *et al*. 2010).

Overall, the MN cluster seems to be more successfully assembled than the LA cluster, which is more fragmented (Figure 5). Differences in assembly completeness between the two regions might be a result of several factors. Firstly, the MN region has more flanking genes in closer proximity to the *Hb* region, which results in a higher density of probes. Secondly, the overall repeat content of the LA region is one order of magnitude larger than in the MN region, largely due to the larger proportion of simple repeats. Repeat content is a major interference in capture experiments because unwanted repetitive DNA may be enriched for, especially if there are repeated sequences included in the probe design. Furthermore, if probes were not completely covered by target DNA they get single-stranded sticky ends that can hybridize to repetitive or other non-target DNA (Newman & Austin 2016). Lastly, unless there were some longer reads that bridged such areas, this would in turn have led to gaps in the downstream *de novo* assemblies. Following that the assembly success was possibly a result of read length, we reason that a future increase of the average read length from 3 kbp to 5-10 kbp, would be sufficient to substantially increase the completeness of the assemblies. Due to the current circular consensus (CCS) PacBio sequencing technology, however, which is a trade-off between accuracy and length of reads, longer reads with sufficient accuracy are not feasible.

### Long-read sequencing capture across species harbors new potential for comparative genomic studies

The number of reads mapping to the target region was in the range of 23-43%, which may be seemingly low compared with other capture studies. For instance, a whole exome capture study on humans reported 56.1% of reads mapped to the target region (Guo *et al*. 2012) and a similar study in rats reported to have 78.3% of reads on target (Yoshihara *et al*. 2016). In contrast to our study however, these capture experiments enriched either the exome or ultra-conserved elements within a single species, allowing for more efficient capture of conserved sequences. We were however, able to cover up to 98% of the target region with a sequencing depth of >10 reads across species (Table S1) which is similar to what mentioned experiments within human and rat exomes reported (Guo *et al*. 2012; Yoshihara *et al*. 2016) and the main difference is the higher percentage of non-target sequences in our study.

We were able to capture complete genes for species with 70 My divergence time from the Atlantic cod (Figure 5). As expected, we found that capture success declines with increased sequence divergence between the reference genome of which we chiefly based our capture probes and the genomes of the included codfishes (Figure 4). It has been reported that orthologous exons were successfully captured in highly divergent frog species (with 200 My of separation), nevertheless the capture success greatly decreased with increased evolutionary distance (Hedtke *et al*. 2013). Similarly, it has been demonstrated that it is possible to capture >97% of orthologous sequences in four species of primates that diverged from humans 40 My ago, using probes entirely based on the human exome (George *et al*. 2011). Further, exomes were effectively captured from skink species that diverged up to 80 My from the reference, yet reporting a substantial decline in capture efficiency for sequences >10 % different from the reference species (Bragg *et al*. 2015). Our study stands out from previous capture experiments because intergenic, noncoding sequences in addition to genes were captured across distantly related species. Efficient capture of intergenic sequences requires less divergence time, as these regions usually evolve faster than genes (Koonin & Wolf 2010). Thus, the most distantly related species from Atlantic cod for which we captured both coding and noncoding sequences was burbot, which diverged from Atlantic cod 46 My (Figure 5). We argue, in line with (Schott *et al*. 2017), that sequence divergence may be a more exact predictor of capture success than evolutionary distance, as the sequence capture process is mainly influenced by the difference between the probe sequence and the target sequence. European hake, marbled moray cod and roughhead grenadier all diverged from cod about 70 My ago, however, the European hake *Hb* regions was more successfully captured and assembled (Table 1; Figure 2). This could be due to European hake having a lower genome-wide divergence to Atlantic cod than marbled moray cod and roughhead grenadier (809k vs 879k and 907k SNPs; Table S1).

Finally, it should be mentioned that cusk – which diverged from Atlantic cod 39 My ago – was added to the experimental design after the probes were generated. Thus, the successful capture of cusk was therefore solely based on cross-species target enrichment, demonstrating the power of heterologous probe targeting.

### Concluding remarks and future perspectives

Here, we have successfully demonstrated that combining targeted sequence capture with long-read sequencing technology is as an efficient approach to obtain high quality sequence data of a specific genomic region, including both coding and noncoding sequences, across evolutionary distant species. We show that genome-wide divergence is of importance for capture success across species. Furthermore, the use of long-read sequencing augmented the *de novo* assembly of regions containing repeated sequences that would otherwise fragment assemblies based on short-read sequencing. This is crucial for capturing complete intergenic sequences that may be highly divergent compared to genic regions even among fairly closely related species. Given the rapid development in sequencing technologies future methods will enable read-through of repeated regions and thus further increase the completeness of assemblies. Moreover, a less stringent hybridization protocol should make it possible to capture sequences across even deeper evolutionary time. In sum, our approach has generated a cross-species genomic resource across distantly related codfishes and show the potential of enhancing comparative genomic studies of continuous genic and intergenic regions between any eukaryotic species-group where genomic resources are scarce.

## Material and methods

### Defining target region and probe design

The probe design was chiefly based on the high-quality genome of Atlantic cod, known as gadMor2 (Tørresen *et al*. 2017). In addition, species-specific probes were designed based on low-coverage assembled genomes (Malmstrøm *et al*. 2016) for ten selected species representing six families in the Gadiformes order. These species were Atlantic cod (*Gadus morhua*), Alaskan Pollock (*Gadus chalcogrammus*), polar cod (*Boreogadus saida*), haddock (*Melanogrammus aeglefinus*), Silvery pout (*Gadiculus argenteus*), burbot (*Lota lota*), European hake (*Merluccius merluccius*), roughhead grenadier (*Macrourus berglax*), roughsnout grenadier (*Trachyrincus scabrus*) and marbled moray cod (*Muraenolepus marmoratus*).

To retrieve relevant sequence data for the probe design, the MN and LA *Hb* regions were extracted from gadMor2 (Figure 1). These sequences, hereby known as the target region, were then used as queries in BLAST (Altschul *et al*. 1990) searches with an E-value threshold of <0.1 against the genome assembly data of all ten species.

In total, 5604 sequences from the chosen species were supplied to NimbleGen probe design. Protein coding genes from the ENSEMBL database were used to define the regions to be tiled in the probe design (Table S5) within the target region of the Atlantic cod, and the unitigs for each of the additional codfishes.

NimbleGen SeqCap EZ capture probes were designed by NimbleGen (Roche, Madison, USA) using a proprietary design algorithm. NimbleGen offers an in-solution sequence capture protocol, which includes custom made probes. Uniquely, the capture probes from NimbleGen are tiled to overlap the target area. 50 – 100 bp (average 75 bp) probes where designed tiled over the target region (subset of gadMor2) resulting in each base, on average, being covered by two probes. Additionally, raw reads from Illumina sequencing from (Malmstrøm *et al*. 2016; 2017) were used for each species to estimate repetitive sequences in each of the species genomes, aiming to discard probes containing any repeats. A more detailed description of the probe design is provided in Supplementary Materials and Methods and Table S6.

### Sample collection and DNA extraction

Our goal working with animals is always to limit any harmful effects of our research on populations and individuals. Whenever possible we try to avoid animals being euthanized to serve our scientific purpose alone by collaborating with commercial fisheries or museums. The tissue samples used in this study are either from commercially fished individuals intended for human consumption or museum specimen. The commercially caught fish were immediately stunned, by bleeding following standard procedures by a local fisherman. There is no specific legislation applicable to this manner of sampling in Norway, however it is in accordance with the guidelines set by the ‘Norwegian consensus platform for replacement, reduction and refinement of animal experiments’ (www.norecopa.no).

DNA was extracted from tissue samples using High Salt DNA Extraction method by Phill Watts (https://www.liverpool.ac.uk/~kempsj/IsolationofDNA.pdf, <u>last day accessed: December 2017</u>). The concentration and purity of the DNA samples were quantified using NanoDrop (Thermo Fisher Scientific, Waltham, MA, USA) and a Qubit fluorometer (Invitrogen, Thermo Fisher Scientific, Waltham, MA, USA). Due to poor DNA quality, three species included in the probe design; Alaskan Pollock, polar cod and roughsnout grenadier were excluded from further analysis. In total, eight species were sequenced; seven of these species were included in the probe design and one closely related species (cusk), which serves as a cross species capture experiment without species-specific probes.

### Capture, library preparation and sequencing

The sequencing libraries were prepared following a modified Pacific Biosciences SeqCap EZ protocol. As multiplexing of the samples before capture was required, barcodes were designed at the Norwegian Sequencing Centre (http://www.sequencing.uio.no) using guidelines from Pacific Biosciences, for more information see Supplementary Materials and methods, Table S7 and Figure S3. Genomic DNA was sheared to 5 kb fragments using MegaRuptor (Diagenode, Seraing (Ougrée), Belgium). Due to poorer DNA quality, fragmenting was not done for European hake. For this sample together with fragmented DNA from roughhead grenadier, short fragments were removed using BluePippin (Sage Science, Beverly, MA, USA) before library preparation. Illumina libraries were prepared using KAPA Hyper Prep kit (Kapa Biosystems, Wilmington, MA, USA) and barcoded using different Illumina barcodes. PacBio barcodes were implemented during pre-capture amplification of libraries. After amplification, fragment length distribution was evaluated using Bioanalyzer (Agilent Technologies, Santa Clara, CA, USA) and samples were pooled in equimolar ratio. During hybridization, SeqCap EZ Developer Reagent (universal repeat blocker for use on vertebrate genomes) and oligos corresponding to Illumina and PacBio barcodes were used for blocking. Captured gDNA was amplified to ensure that sufficient amount of DNA was available for PacBio library preparation. Size selection of the libraries was performed using BluePippin. Final libraries were quality checked using Bioanalyzer and Qubit fluorometer (Invitrogen, Thermo Fisher Scientific, Waltham, MA, USA) and sequenced on RS II instrument (PacBio, Menlo Park, CA, USA) using P6-C4 chemistry with 360 minutes movie time. In total, 9 SMRT cells were used for sequencing.

### *De novo* assemblies

Reads were filtered and de-multiplexed using the ‘RS_reads of insert.1’ pipeline on SMRT Portal (SMRT Analysis version smrtanalysis_2.3.0.140936.p2.144836). Each set of reads corresponding to a given species was crossed-checked with their respective six-nucleotide Illumina adapter. Reads containing an incorrect Illumina adapter were removed. Adapter sequences were then trimmed using the application Prinseq-lite v0.20.4 (Schmieder & Edwards 2011). The trimmed reads were assembled *de novo* using Canu v1.4 +155 changes (r8150 c0a988b6a106c27c6f993dfe586d2336282336a6) (Berlin *et al*. 2015). The Canu software is optimized for assembling single molecule high noise sequence data. We specified genome size as the size of the target region (300 kbp). Additionally, we ran PBJelly (English *et al*. 2012) on the Canu *de novo* assemblies, using the raw reads to possible bridge gaps between scaffolds, settings given in Supplementary Materials and Methods.

We assessed the assemblies by running Assemblathon 2 (Bradnam *et al*. 2013), which reports assembly metrics such as the longest contig, the number of contigs, and the N50 value. *De novo* assemblies of the MN and LA regions of Atlantic cod and haddock were aligned and compared to their reference genomes, gadMor2 and melAeg respectively, using BLAST and BWA v0.7.10 (Li & Durbin 2009) to determine syntenic similarities and assembly completeness.

### Estimating capture success

PacBio reads for all the species were mapped back to the Atlantic cod genome assembly (gadMor2) in order to determine sequence capture success and target mapping depths. Mapping was done using BWA-MEM v0.7.10 (Li & Durbin 2009). Target-area read depth for all the species based on mapping against gadMor2, were calculated using Samtools v1.3.1 (Li *et al*. 2009). We calculated both average and median mapping depth against the target region as a whole and for the MN and LA region separately. We also calculated percentage of reads that mapped to the target region, and the percentage of the target regions covered by reads to a minimum depth of 10x. To compare assemblies to the target region we additionally mapped the assemblies to the target region. In order to verify the sequence capture process, sequence data for Atlantic cod and haddock were mapped back to their reference genomes using BWA-mem v0.7.10 (Li & Durbin 2009). The results were visualized using Integrative Genome Viewer (Robinson *et al*. 2011).

To obtain an independent measure of divergence between species in the capture experiment we calculated genome wide level of divergence of each species to the reference genome of Atlantic cod using low-coverage whole-genome sequence data from (Malmstrøm *et al*. 2016; 2017). We mapped raw reads to Atlantic cod using BWA-MEM (Li & Durbin 2009) and called SNPs using the Freebayes variant caller (Garrison & Marth 2012). Some species are more closely related to Atlantic cod than others, which could introduce a bias in mapping. To avoid this, we only looked at genomic regions where all species mapped. The number of SNPs was then used as an estimate of genome-wide divergence of each species to Atlantic cod. We also mapped a low-coverage genome of Atlantic cod to the Atlantic cod reference genome as a control.

In pursuance of factors explaining capture success we tested for correlations and plotted the relationship between the genome wide level of divergence and the following variables; median mapping depth against the target region (for total, LA and MN, respectively); percentage of reads that mapped to the target region; and the percentage of the target region covered by reads. All tests and plots were done using R version 3.2.5 (R Core Team 2016).

Assembly continuity is very often hampered by the presence of repeats, which create gaps. We therefore quantified repeat-content in the target region extracted from gadMor2 and orthologous regions in haddock using Repeatmasker Open 3.0 (Smit A *et al*. 2010) for the MN region and the LA region separately.

### Identifying gene location and synteny

In order to identify the genes of interest and their location in the assembly we used local sequence alignment algorithm BLAST v2.4.0 (Altschul *et al*. 1990) with protein sequences of the genes of interest (Table S5) as queries. tblastn was used with an e-value of 0.1. Investigation of *Hbb1-Hba1* promoter region was done for four species, Atlantic cod, haddock, silvery cod and cusk. Sequences were aligned with ClustalW default settings using MEGA7 (Kumar *et al*. 2016). Ancestral sequence reconstruction was carried out for *Hbb-1* gene sequences from 24 species of codfishes from (Baalsrud *et al*. 2017) using a maximum likelihood method implemented in MEGA7 (Kumar *et al*. 2016).

Additionally, we estimated sequence identity using EMBOSS Needle (Rice *et al*. 2000) with default settings, between *Hbb* gene sequences from (Baalsrud *et al*. 2017) that were missing and present in the *de novo* assemblies to evaluate similarity (Table S2).

## Acknowledgements

We would like to thank Marianne H. Selander Hansen and Alexander J. Nederbragt for help with the initial design of this project. All computational work was performed on the Abel Supercomputing Cluster (Norwegian Metacenter for High-Performance Computing (NOTUR) and the University of Oslo), operated by the Research Computing Services group at USIT, the University of Oslo IT Department. Sequencing library creation and high-throughput sequencing were carried out at the Norwegian Sequencing Centre (NSC), University of Oslo, Norway. This work was funded by a grant awarded to K.S.J. from the Research Council of Norway (RCN grant 222378).

## Author contributions

H.T.B. and S.J. initially conceived and designed the study, with input from S.N.K.H, A.T.-K., M.S., G.O., R.S., and K.S.J. Tissue samples were provided by S.J. and H.T.B. Probe design was carried out by T.R. with assistance from S.N.K.H and H.T.B. DNA extraction and sequence library preparation was performed by S.N.K.H and A.T.-K, respectively. Sequence capture was carried out by S.N.K.H, A.T.-K., M.S. and G.O. Filtering, mapping of sequences and *de novo* assemblies was done by S.N.K.H., assisted by O.K.T and H.T.B. Annotation of genes, synteny analyses, statistical analyses and construction of all figures and tables was done by S.N.K.H and H.T.B. The manuscript was written by S.N.K.H and H.T.B. with input from S.J. and K.S.J.

## Competing interests

The authors declare that they have no competing interests.

## Data and materials availability

All reads and assemblies (unitigs) reported on here, and the target region, subset of gadMor2, as well as the relevant sequence data for the probe design from the chosen species supplied to NimbleGen and the probe sequences have been deposited at figshare under doi/xxx.

